# Population dynamics and transcriptomic responses of *Pseudomonas aeruginosa* in a complex laboratory microbial community

**DOI:** 10.1101/351668

**Authors:** Yingying Cheng, Joey Kuok Hoong Yam, Zhao Cai, Yichen Ding, Lian-Hui Zhang, Yinyue Deng, Liang Yang

## Abstract

*Pseudomonas aeruginosa* is one of the dominant species when it co-exists with many other bacterial species in diverse environments. To understand its physiology and interactions with co-existing bacterial species in different conditions, we established physiologically reproducible eighteen-species communities, and found that *P. aeruginosa* became the dominant species in mixed-species biofilm community but not in the planktonic community. *P. aeruginosa* H1 type VI secretion system was highly induced in the mixed-species biofilm community compare to its mono-species biofilm, which was further demonstrated to play a key role for *P. aeruginosa* to gain fitness over other bacterial species. In addition, the type IV pili and Psl exopolysaccharide were shown to be required for *P. aeruginosa* to compete with other bacterial species in the biofilm community. Our study showed that the physiology of *P. aeruginosa* is strongly affected by interspecies interactions, and both biofilm determinants and H1 type VI secretion system contribute to *P. aeruginosa* fitness over other species in complex biofilm communities.

**Importance:** *Pseudomonas aeruginosa* usually coexists with different bacterial species in natural environment. However, systematic comparative characterization of *P. aeruginosa* in complex microbial communities with its mono-species communities is lacking. We constructed mixed-species planktonic and biofilm communities consisting *P. aeruginosa* and seventeen other bacterial species to study the physiology and interaction of *P. aeruginosa* in complex multiple-species community. A single molecule detection platform, NanoString nCounter^®^ 16S rRNA array, was used to shown that *P. aeruginosa* can become the dominant species in the biofilm communities while not in the planktonic communities. Comparative transcriptomic analysis and fluorescence-based quantification further revealed that *P. aeruginosa* H1 type VI secretion system and biofilm determinants are both required for its fitness in mixed-species biofilm communities.

## Introduction

Biofilm can protect bacteria from hostile environment and also disperse bacteria to colonize in new niches (1, 2). *Pseudomonas aeruginosa* is a model organism for studying biofilm formation in Gram-negative bacteria (1, 3). Type IV pili (T4P), flagellar, iron uptake system, extracellular DNA (eDNA) and Pel/Psl exopolysaccharides (EPS) are well known factors that contribute to the development of *P. aeruginosa* mono-species biofilm (4–6). The biofilm of *P. aeruginosa* makes it more difficult to treat (7, 8), thus expands its survival range in nature. The adaptability of *P. aeruginosa* to various environments, such as water, soil, sewage, and plants, suggests its strong competitiveness for ecological niches (4, 9, 10). For example, *P. aeruginosa* is a dominant cultivable endophytic bacterium associated with *Pennisetum glaucum* (11).

The broad spectrum adaptability of *P. aeruginosa* also hints its strong competitiveness against various other bacterial species when in mixed-species microbial communities. Several studies indicated that *P. aeruginosa* is able to gain fitness over other species during its survival in either two-species or three-species co-cultures (12–14). For example, *P. aeruginosa* reduces the viability of *Staphylococcus aureus* and lyses *S. aureus* cells for iron repletion in planktonic co-cultures (15, 16). In biofilms, *P. aeruginosa* inhibits *S. epidermidis* growth and reduces the initial adhesion of *Agrobacterium tumefaciens* through the secretion of Psl and Pel exopolysaccharides and small diffusible molecules (17–19). *P. aeruginosa* was shown to decrease the swarming motility of *P. putida* and therefore inhibit its biofilm formation by producing 2-heptyl-3-hydroxy-4-quinolone (20). Quinolones secreted by *P. aeruginosa* were also able to reduce biofilm maturation for a broad range of bacteria (21). These studies proved the competitiveness of *P. aeruginosa* in mixed-species interaction results from metabolic versatility and strong biofilm formation capacity. The known devices for *P. aeruginosa* to implement competition including type VI secretion system (T6SS), antibiotic, iron chelators, pyoverdine, rhamnolipid, pyocyanin, extracellular polysaccharide, fatty acid cis-2-decenoic acid, proteinase and other quorum sensing system regulated virulence (22–27).

*P. aeruginosa* co-exists with many different microbial species in actual environment. However, the main competition advantage of *P. aeruginosa* in gaining fitness over other species within complex microbial communities remained unclear. Previous studies indicated that interspecies interactions in mixed-species microbial communities are complicated and involve both cooperation and competition (24, 28). Therefore, monitoring the population dynamics of *P. aeruginosa* in complex microbial communities is important for elucidating how this interaction network is established, which may provide insights into *P. aeruginosa* ecological role in complex mix-species communities.

Until now, there is a lack of robust tool to monitor the population dynamics in complex microbial communities. The high throughput digital NanoString nCounter® system has recently been used to profile the multiplexed gene expression with flexibility and sensitivity (29, 30). Here we evaluated this technology in monitoring population dynamics in a laboratory eighteen-species microbial community with current platform. We further investigated the transcriptome of *P. aeruginosa* competition in this mixed-species community in both planktonic and biofilm modes of growth.

## Results

### Establishment of the complex microbial community

Instead of proliferating as a single species culture, *P. aeruginosa* often grows as common species within mixed-species microbial communities containing many other bacterial species in natural environments as well as infection sites (31, 32). We presumed bacteria with same region colonization have chance to co-exist. To obtain a better understanding of *P. aeruginosa* physiology in complex microbial communities and evaluate the NanoString nCounter® system, we selected bacterial species that potentially co-exist with *P. aeruginosa* in different environments and can be distinguished from each other by NanoString nCountern ® probes, including both of human pathogens and environmental bacteria (Supplementary Table 1), to establish laboratorial planktonic and biofilm communities. *Acinetobacter baumannii, Burkholderia cenocepacia, Klebsiella pneumoniae* and *S. aureus* can co-infect lungs with *P. aeruginosa* (33–36); *Citrobacter amalonaticus, E. coli* and *Streptococcus gallolyticus* can cause intra-abdominal infections like *P. aeruginosa*; *Chromobacterium violaceum, S. aureus* and *Stenotrophomonas maltophilia* can infect the skin or open wounds as *P. aeruginosa* can; *Listeria monocytogenes* may colonize in host from gastrointestinal tracts as *P. aeruginosa; P. putida* can be isolated from infected urinary tract and infected skin as well as *P. aeruginosa; Bacillus subtilis, Elizabethkingia meningoseptica, P. fluorescens* and *Shewanella oneidensis* are widely distributed in natural environments as *P. aeruginosa. P. aeruginosa* was also isolated from plant as *P. syringae* and *Xanthomonas campestris* (37). Both of known and undiscovered natural environments are so complex and unexpected. This laboratory community may help us to understand *P. aeruginosa* real response in possible complex microbial community. Not only to investigate the ecological behaviour and physiological response of *P. aeruginosa* in our model system; we also hope to provide insights into how future studies on complex microbial communities can be shaped. We used 10% Tryptic Soy Broth (TSB) as the growth medium, which is able to support the growth of all the selected eighteen bacterial species at room temperature (Supplementary Figure 1). Although *A. baumannii, C. amalonaticus, C. violaceum, E. meningoseptica, E. coli* and *K. pneumoniae* grew relatively faster than other species, whereas *L. monocytogenes* and *S. gallolyticus* were unable to grow to a high cell density in 10% TSB, the growth curves indicated that 10% TSB is generally suitable for establishing the mixed-species microbial community for *P. aeruginosa* to grow at a comparable rate compared to these species (Supplementary Figure 1).

In addition to the growth rate, the biofilm formation capacity of these bacterial species was also tested. After 24 hours of static incubation, A. *baumannii* and *K. pneumoniae* had the highest capacity to form biofilms (Supplementary Figure 2). Among the other sixteen species, *B. cenocepacia, P. aeruginosa* and *S. maltophilia* have similar biofilm formation capabilities (Supplementary Figure 2). However, *B. subtilis, C violaceum, E. meningoseptica, P. putida, P. syringae* and *S. gallolyticus* formed less biofilms in 10% TSB at room temperature (Supplementary Figure 2). In summary, the biofilm formation capacities of these eighteen species are rather different, with no linear relationship between their biofilm formation capacity and growth rates. We therefore established the planktonic and biofilm microbial communities by adjusting each species to an OD_600_ value of 0.01 in 10% TSB medium as input.

### Physiological reproducibility of the mixed-species microbial communities

Before investigating the population dynamics and physiology of *P. aeruginosa* in the mixed-species communities, we employed RNA sequencing-based metatranscriptomics to examine the physiological reproducibility of both the planktonic and biofilm communities. The sequencing reads (7.8–9.2 million per sample) were assigned to 44 classified functional roles (SEED; 51.4 million reads) with unclassified reads in MEGAN6. Similar profiles of functional classifications of genes assigned to each community model by their functional properties were observed among three biological replicates (Figure 1A), indicating that both planktonic and biofilm community physiology are highly reproducible in our laboratory model (Figure 1A). In addition, the principal coordinates analysis (PCoA) confirmed that the metatranscriptome profiles of planktonic community are different from biofilm community, and the three biological replicates for each community were similar because they clustered in close proximity (Figure 1B), which suggests that the biological replicates of the mixed-species cultures have a reproducible physiology and be able to serve as functional communities in our models.

**Figure 1.**
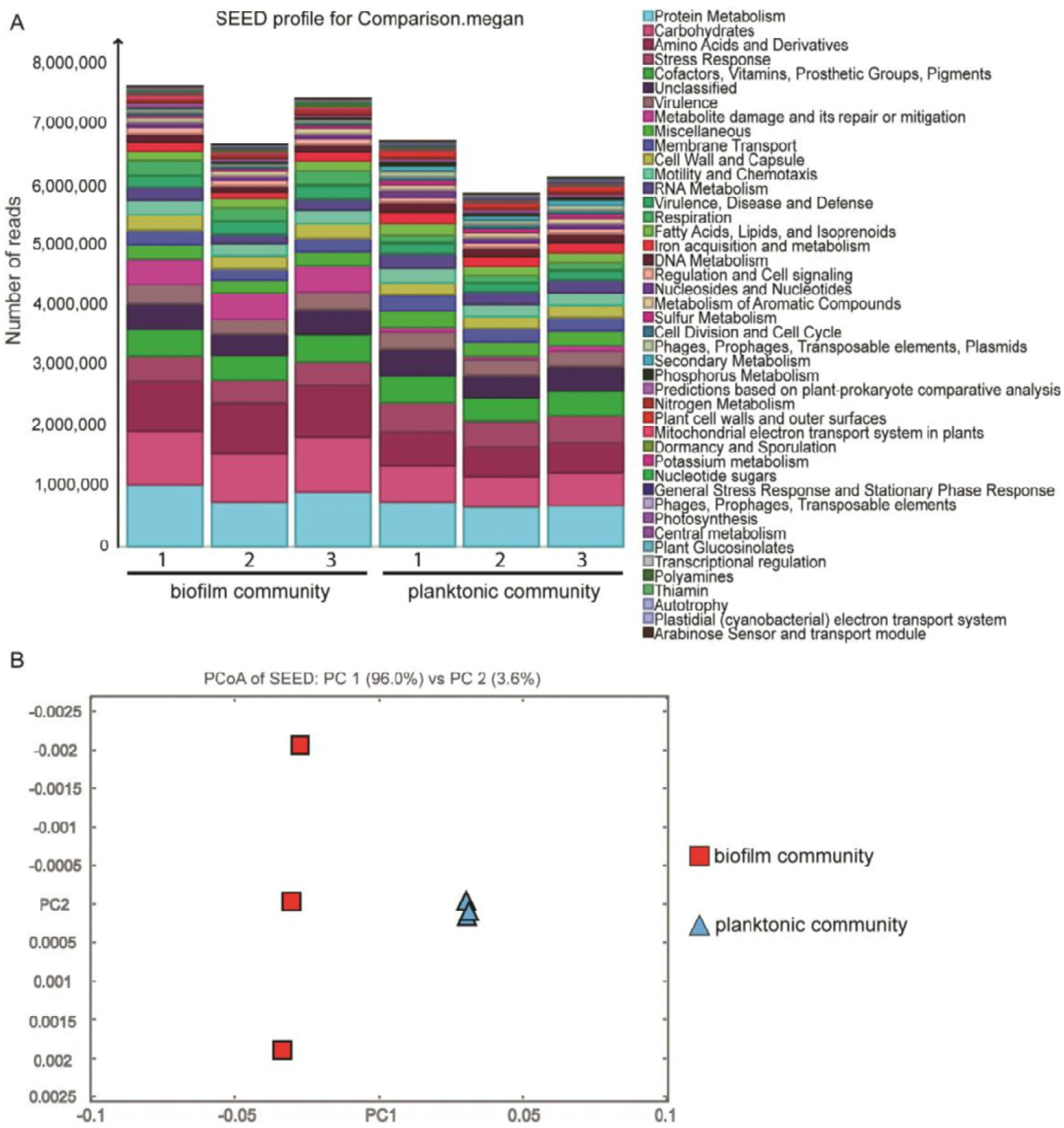
Metatranscriptomics comparison between mixed-species planktonic community and biofilm community modes. A) Functional classification by SEED. The comparison was based on read counts with a minimum quality score cut-off of 50 and a top percentage of 25. Different colours corresponding to different functions are ranked in the right column. B) Clustering by PCoA based on SEED classification using Bray-Curtis dissimilarity.

### Evaluation of NanoString nCounter® 16S rRNA array for microbial population assay

Although the 16S rRNA gene amplicon sequencing approach is widely used to study the microbial population diversity, it is known to have the drawback of PCR bias. A NanoString nCounter® 16S rRNA array was developed to detect unique signals from complex hybridized samples at a single molecule level by utilizing special probes and omitting the PCR amplification step (38) (Supplementary Table 1). To check the feasibility of the designed nCounter® 16S rRNA array for detecting the 16S rRNA from different bacterial species in the total RNA mixture, we mixed the total RNA extracted from fourteen individual bacterial species at different ratios in three different training groups (Table 1) and subjected them to nCounter® 16S rRNA array analysis with the total RNA from the planktonic and biofilm communities together. The other three *Pseudomonas* species were not mixed in the training groups to test the background of *Pseudomonas* probe to other species in the mixed-species community by applying this new technology. *L. monicytogenes* as gram-positive representation was not mixed to test the gram-positive bacterial background. The percentage of each species in 16S rRNA reads according to nCounter® 16S rRNA array analysis was compared to its ‘actual’ RNA percentage from the training groups to calculate the normalization factors for each of the fourteen species (Table 1). Table 1 shows that the nCounter® 16S rRNA array gave some species more reads than accurate 16S rRNA readings, while some other species got less reads compare to true 16S rRNA readings. This trend indicated that it is crucial to analyse the bacterial population dynamics by multiplying the normalization factors (the variation fold) (Table 1), when nCounter® 16S rRNA array is employed to reflect the true percentage of each bacterial species in a mixed-species community.

**Table 1.** 16S rRNA sequencing feasibility detection. Three different random mixed groups were named “I”, “II” and “III” in this table. Numbers indicate the ratio of one species in the mixed group with unit %. “Reads percentage” is 16S rRNA sequencing analysis result; “RNA percentage” shows proportion of single species total RNA in mixture. “Fold” calculated from average values of ratio in Reads percentage to it in RNA percentage. Dash in RNA percentage means no RNA was mixed into the random mixture sample, so some of average fold values and standard deviations cannot been calculated, also labelled with dash.

|  | Reads percentage (%) |  |  | RNA percentage (%) |  |  | Fold (normalization factor) |
| --- | --- | --- | --- | --- | --- | --- | --- |
|  | I | II | III | I | II | III |  |
| *A. baumannii* | 3.13 | 6.52 | 1.09 | 1.95 | 3.71 | 1.92 | 1.31 ±0.65 |
| *B. subtilis* | 0.18 | 0.07 | 0.06 | 0.11 | 0.10 | 0.10 | 1.00 ±0.61 |
| *B. cenocepacia* | 3.88 | 0.54 | 1.01 | 1.93 | 1.76 | 9.55 | 0.81 ±1.04 |
| *C. violaceum* | 16.94 | 3.43 | 3.05 | 21.48 | 9.77 | 10.60 | 0.48 ±0.27 |
| *C. amalonaticus* | 7.56 | 6.36 | 0.10 | 11.45 | 20.84 | - | 0.48 ±0.25 |
| *E. coli* | 2.10 | 2.50 | 0.97 | 1.88 | 6.83 | 3.71 | 0.58 ±0.47 |
| *E. anophelis* | 0.87 | 0.00 | 0.00 | 0.41 | - | - | 2.10 ±- |
| *K. pneumoniae* | 6.90 | 0.29 | 0.11 | 40.48 | - | - | 0.17 ±- |
| *L. monocytogenes* | 0.01 | 0.02 | 0.03 | - | - | - | - |
| *P. aeruginosa* | 23.28 | 47.95 | 72.63 | 6.53 | 29.69 | 57.95 | 2.14 ±1.24 |
| *P. fluorescens* | 0.44 | 0.92 | 1.46 | - | - | - | - |
| *P. putida* | 2.22 | 4.98 | 6.13 | - | - | - | - |
| *P. syringae* | 0.01 | 0.01 | 0.01 | - | - | - | - |
| *S. oneidensis* | 0.28 | 0.35 | 0.24 | 1.24 | 2.25 | 2.44 | 0.16 ±0.06 |
| *S. aureus* | 13.24 | 5.18 | 2.09 | 3.73 | 3.40 | 1.84 | 2.07 ±1.30 |
| *S. maltophilia* | 0.47 | 0.01 | 0.00 | 0.59 | - | - | 0.79 ±- |
| *S. gallolyticus* | 2.67 | 10.34 | 0.91 | 0.58 | 4.25 | 0.58 | 2.86 ±1.54 |
| *X. campestris* | 15.82 | 10.52 | 10.10 | 7.65 | 17.40 | 11.32 | 1.19 ±0.78 |

### *P. aeruginosa* is the dominant species in the mixed-species biofilm community

In order to maintain the reproducibility of this laboratory model, we still use the same mixed-species community. Each individual species of bacterial cells was normalized to an OD_600_ reading of 0.01 in the mixture and cultivated for 1 day to establish the planktonic community, and 5 days to establish the biofilm community, and total RNA extracted from these communities were subjected to analysis by nCounter® 16S rRNA array. In the 1-day-old planktonic community, the relative abundance of *A. baumannii, C. violaceum* and *E. meningoseptica* increased by 2-fold or more compared to their initial inputs, whereas the relative abundance of *B. subtilis*, *P. aeruginosa*, *S. oneidensis*, *S. aureus*, *S. gallolyticus* and *X. campestris* decreased to half or less compared to their initial inputs (Figure 2 and Supplementary Table 2). Interestingly, the microbial population dynamics are quite different in the 5-day-old biofilm community compared to the planktonic community; the relative abundance of *K. pneumoniae* and *P. aeruginosa* increased to more than 4-fold their initial inputs, whereas most of the other species decreased significantly (Figure 2 and Supplementary Table 2). *P. aeruginosa* 16S rRNA constituted around 32.62% of the total 16S rRNA content while *K. pneumoniae* constituted around 26.08% of that in 5-day-old biofilms (Figure 2, and Supplementary Table 2) after adjusted by normalization factors (Table 1). The proportions of all bacterial species (including the four non-normalized ones) are summarized in Supplementary Figure 3.

**Figure 2.**
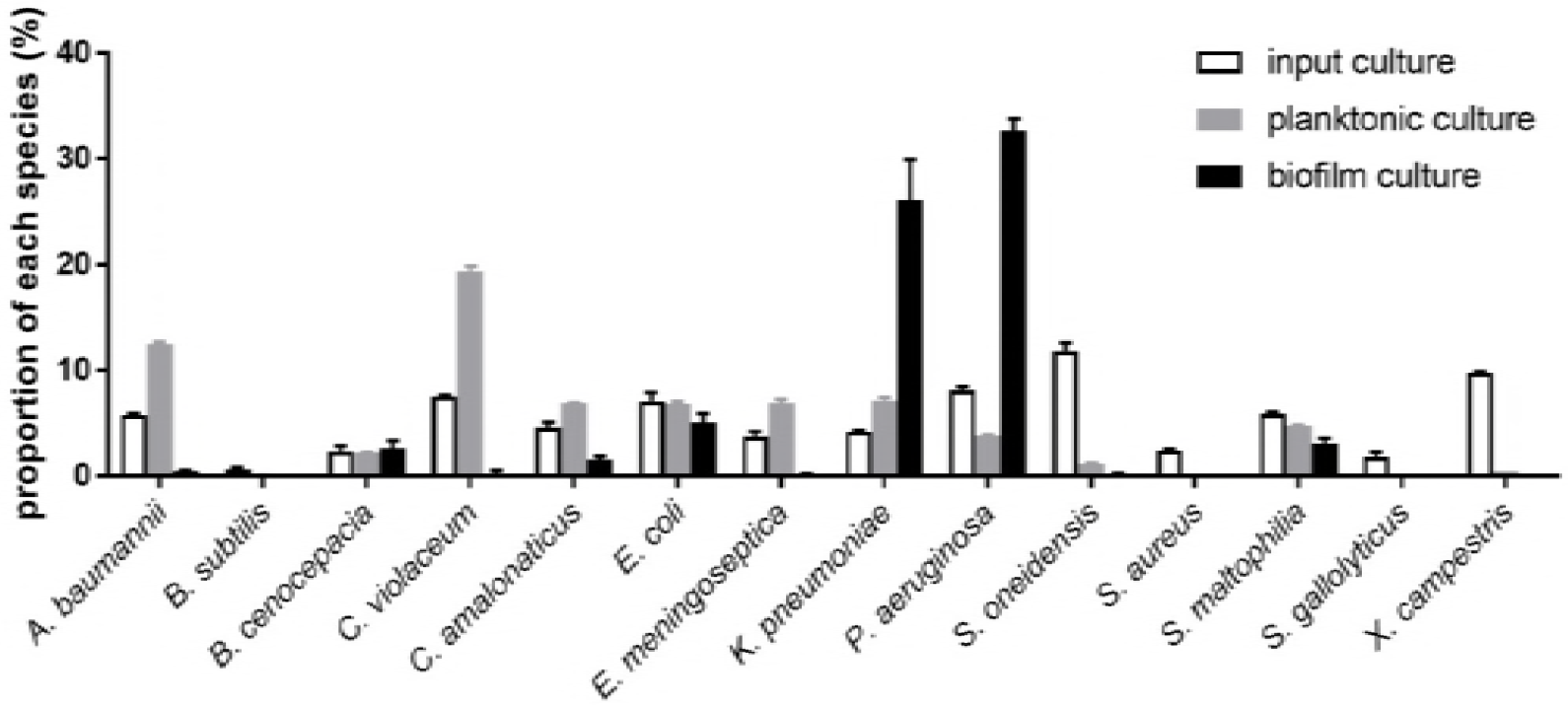
Normalized population dynamics of each species in the mixed-species microbial communities. The input culture is the mixture of all species mixed at equal OD_600_ amounts at the starting point for both planktonic and biofilm incubation. Grey bars indicate the proportion of each species in the 24 h-incubated planktonic microbial community. The black bars show the population dynamics of each species in the 5-day biofilm microbial community.

To validate the observed microbial population dynamics, we used GFP-tagged *P. aeruginosa* to establish the 1-day-old planktonic community and the 5-day-old biofilm community as described above. The abundance of *P. aeruginosa* was detected and calculated using the fluorescence-proportion standard curve (Supplementary Figure 4). The fluorescence-proportion standard curve is preferred over the CFU counting approach due to the difference in the growth rates and the colony sizes of these eighteen bacterial species on plates, which compromises the accuracy of CFU counting. Figure 3A shows that the results of the biological fluorescence-proportion standard curve-based experiment correlated well with the results of the normalized nCounter® 16S rRNA array. Cells from these fluorescence-based proportion tests were resuspended and examined using confocal microscopy to verify that the proportions of *P. aeruginosa* cells in the mixed-species biofilm community (Figure 3C) is much higher than in the planktonic community (Figure 3B). Together with the results in Figure 2, we have demonstrated that *P. aeruginosa* is the dominant species in this microbial biofilm community.

**Figure 3.**
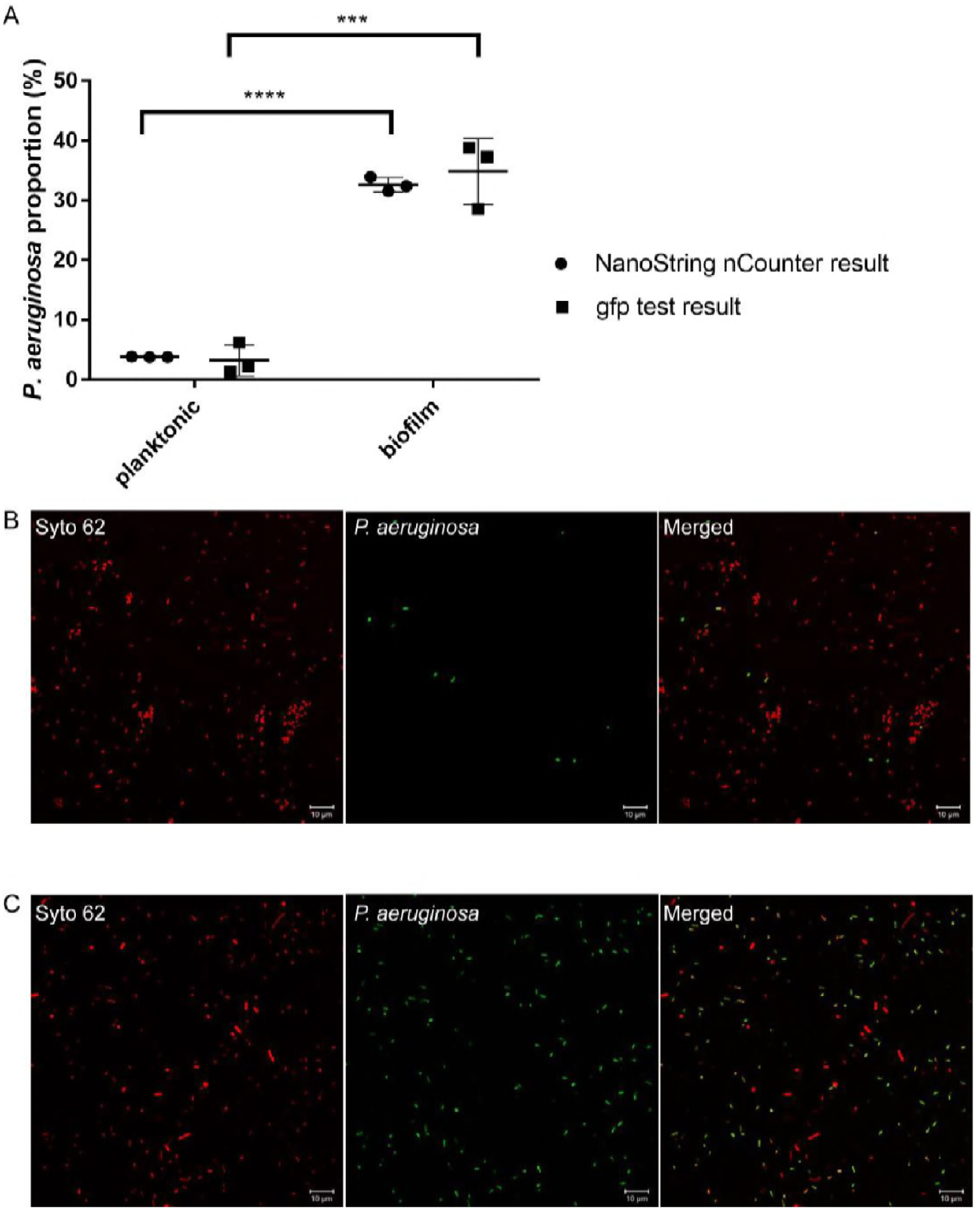
Proportions of *P. aeruginosa* in the mixed-species microbial communities. A) Circular dots indicate three biological replicates of *P. aeruginosa* normalized proportions in both the planktonic and biofilm communities as measured by NanoString nCounter® 16S rRNA array; the square dots show three biological replicates of *P. aeruginosa* proportions in mixed-species microbial communities as calculated by fluorescence-based proportion test. The lines show the average values and standard deviation. Comparison was performed by t-test with p<0.05. B) Confocal images of the resuspended 24 h-incubated mixed-species planktonic microbial community containing GFP-tagged *P. aeruginosa.* C) Confocal images of the resuspended 5-day mixed-species biofilm community containing GFP-tagged *P. aeruginosa.* Green cells show the GFP-tagged *P. aeruginosa,* red cells show all the living cells stained with Syto62.

### Both the H1 Type VI secretion system and biofilm formation determinants contributed to the fitness of *P. aeruginosa* in the biofilm community

Since *P. aeruginosa* constituted up to 32.62% of the biofilm community in contrast to only 3.81% in the planktonic community (Supplementary Table 2), we hypothesized that the important biofilm determinants are likely to contribute to the dominating competence of *P. aeruginosa* over other species in the biofilm community. To test this hypothesis, we performed a transcriptomics analysis to compare the *P. aeruginosa* gene expressions in mono-species biofilms and in the mixed-species biofilm community. Differentially expressed genes of *P. aeruginosa* in the mixed-species biofilm and mono-species biofilm are illustrated as a heat map (Figure 4A). Further clustering of the transcriptome by Principle Component Analysis (PCA) revealed that the *P. aeruginosa* maintained a distinct physiology in these two types of biofilms (Figure 4B). Based on the negative binomial test with adjusted p-value cut-off of 0.05 and a log_2_ fold-change cut-off of 1, we found that 239 genes were upregulated and 171 genes were downregulated in *P. aeruginosa* cells in the mixed-species biofilm compared to those in mono-species biofilms (Supplementary Table 3).

Among the significantly upregulated genes of *P. aeruginosa* in the mixed-species biofilm, at least 36 genes belong to the T6SS (Table 2), which accounted for 15% of all the upregulated genes. Most of them belong to Hcp secretion island I (H1) T6SS while some are located in the Hcp secretion island II (H2) T6SS. However, there are only a few genes that belong to H3-T6SS. The ClpV protein is the energy source of T6SS (39); the expressions of *clpV1, clpV2* and *clpV3* belonging to three different sub-type T6SS in *P. aeruginosa* were all upregulated in the mixed-species biofilm (Table 2). To validate the transcriptomic analysis results, the Δ*clpV1*, Δ*clpV2* and Δ*clpV3* mutants of *P. aeruginosa* were constructed with a GFP tag and thereafter formed biofilms with the other seventeen species. Interestingly, only the Δ*clpV1* mutant was greatly impaired in terms of fitness to dominate in the mixed-species biofilms; when ClpV1 was complemented into this mutant with multiple copies, the complementary strain showed higher fitness to dominate in the mixed-species biofilm community than both wild-type PAO1 and the deletion mutant (Figure 5). This finding suggests that the H1-T6SS has a profound effect on fitness gains for *P. aeruginosa* to outcompete other species in the mixed biofilm community (Figure 5). H1-T6SS contains seven effectors to be secreted into its competitor, which are termed Tse1-7 (40, 41). Both Tse1 and Tse3, which can dissolve the target cells by degrading their peptidoglycan (26), were upregulated in the complex microbial biofilm community (Table 2). Furthermore, the expression of the gene encoding the H1-T6SS delivery-dependent proteins VgrG1 was also increased (Table 2).

**Table 2.** Part of *P. aeruginosa* regulated genes in mixed-species biofilm community.

| Locus tag | Gene name | Fold change | p value | Product | Pathway |
| --- | --- | --- | --- | --- | --- |
| PA0070 | *tagQ1* | 2.66 | 1.20E-09 | TagQ1 | H1-T6SS |
| PA0074 | *ppkA* | 2.06 | 1.77E-04 | serine/threonine protein kinase PpkA | H1-T6SS |
| PA0077 | *icmF1* | 2.43 | 2.68E-07 | IcmF1 | H1-T6SS |
| PA0079 | *tssK1* | 2.06 | 4.11E-04 | TssK1 | H1-T6SS |
| PA0082 | *tssA1* | 2.27 | 5.28E-04 | TssA1 | H1-T6SS |
| PA0083 | *tssB1* | 5.34 | 1.81E-23 | TssB1 | H1-T6SS |
| PA0084 | *tssC1* | 5.24 | 1.51E-31 | TssC1 | H1-T6SS |
| PA0085 | *hcp1* | 4.82 | 1.20E-52 | Hcp1 | H1-T6SS |
| PA0086 | *tagJ1* | 2.43 | 4.84E-04 | TagJ1 | H1-T6SS |
| PA0088 | *tssF1* | 2.85 | 6.79E-07 | TssF1 | H1-T6SS |
| PA0089 | *tssG1* | 2.62 | 8.73E-04 | TssG1 | H1-T6SS |
| PA0090 | *clpV1* | 4.96 | 1.52E-21 | ClpV1 | H1-T6SS |
| PA0091 | *vgrG1* | 3.53 | 1.57E-12 | VgrG1 | H1-T6SS |
| PA0092 | *tsi6* | 2.4 | 8.41E-04 | Tsi6 | H1-T6SS |
| PA0097 | PA0097 | 2.48 | 7.21E-05 | hypothetical protein |  |
| PA0099 | PA0099 | 3.61 | 2.72E-15 | type VI effector protein |  |
| PA0100 | PA0100 | 2.86 | 8.24E-15 | hypothetical protein |  |
| PA0260 | *tle3* | 2.62 | 2.67E-17 | Tle3 | H2-T6SS |
| PA0261 | PA0261 | 2.34 | 2.26E-05 | hypothetical protein |  |
| PA0263 | *hcpC* | 4.23 | 1.31E-18 | secreted protein Hcp |  |
| PA1509 | *tplEi* | 2.12 | 1.83E-07 | immunity protein TplEi |  |
| PA1510 | *tplE* | 2.8 | 1.54E-12 | type 6 PGAP1-like effector, TplE | H2-T6SS |
| PA1512 | *hcpA* | 3.52 | 1.95E-14 | secreted protein Hcp |  |
| PA1657 | *tssB2* | 3.06 | 7.18E-18 | TssB2 | H2-T6SS |
| PA1658 | *tssC2* | 2.65 | 1.06E-16 | TssC2 | H2-T6SS |
| PA1659 | *tssE2* | 2.18 | 8.21E-05 | TssE2 | H2-T6SS |
| PA1661 | *tssH2* | 2.35 | 1.57E-06 | TssH2 | H2-T6SS |
| PA1662 | *clpV2* | 2.36 | 1.79E-10 | clpV2 | H2-T6SS |
| PA1663 | *sfa2* | 2.61 | 5.56E-08 | Sfa2 | H2-T6SS |
| PA1665 | *fha2* | 2.74 | 3.11E-09 | Fha2 | H2-T6SS |
| PA1666 | *lip2* | 3.3 | 2.74E-11 | Lip2 | H2-T6SS |
| PA1667 | *tssJ2* | 2.8 | 5.26E-08 | TssJ2 | H2-T6SS |
| PA1668 | *dotU2* | 2.27 | 1.18E-05 | DotU2 | H2-T6SS |
| PA1669 | *icmF2* | 3.21 | 1.74E-14 | IcmF2 | H2-T6SS |
| PA1844 | *tse1* | 2.86 | 1.62E-03 | Tse1 | H1-T6SS |
| PA1845 | *tsi1* | 2.81 | 4.93E-04 | Tsi1 | H1-T6SS |
| PA2365 | *tssB3* | 3.78 | 6.97E-22 | TssB3 | H3-T6SS |
| PA2366 | *tssC3* | 3.75 | 1.81E-37 | TssC3 | H3-T6SS |
| PA2367 | *hcp3* | 3.08 | 1.35E-18 | Hcp3 | H3-T6SS |
| PA2369 | *tssG3* | 2 | 1.03E-04 | TssG3 | H3-T6SS |
| PA2371 | *clpV3* | 2.11 | 1.07E-08 | ClpV3 | H3-T6SS |
| PA3294 | *vgrG4a* | 2.17 | 1.02E-03 | VgrG4a | H2-T6SS |
| PA3484 | *tse3* | 2.54 | 6.03E-04 | Tse3 | H1-T6SS |
| PA3479 | *rhlA* | -2.35 | 4.99E-07 | rhamnosyltransferase chain A | Quorum sensing |
| PA3478 | *rhlB* | -2.17 | 8.81E-07 | rhamnosyltransferase chain B | Quorum sensing |
| PA1871 | *lasA* | -2.42 | 1.64E-08 | LasA protease precursor | Quorum sensing |
| PA3724 | *lasB* | -2.72 | 3.03E-12 | elastase LasB | Quorum sensing |
| PA1432 | *lasI* | 2.43 | 1.13E-14 | autoinducer synthesis protein LasI | Quorum sensing |
| PA2587 | *pqsH* | -2.09 | 3.42E-05 | probable FAD-dependent monooxygenase | Quorum sensing |
| PA1078 | *flgC* | 2.08 | 1.14E-05 | flagellar basal-body rod protein FlgC | Flagella assembly |
| PA1079 | *flgD* | 2.26 | 2.60E-10 | flagellar basal-body rod modification protein FlgD | Flagella assembly |
| PA3059 | *pelF* | 2.44 | 2.40E-03 | PelF | extracellular polysaccharide<br>biosynthesis |
| PA3058 | *pelG* | 2.82 | 1.84E-03 | PelG | extracellular polysaccharide<br>biosynthesis |

In addition to T6SS, the *P. aeruginosa pelF, pelG, flgC* and *flgD* genes, which are important for its biofilm formation in mono-species cultures, were also upregulated in the mixed-species biofilms (Table 2). We next investigated the impact of the classic mono-species *P. aeruginosa* biofilm determinants, such as the Pel and Psl EPS (42–44), T4P (6, 45) and flagella (6, 45), on the fitness of *P. aeruginosa* in the mixed-species biofilms. The *P. aeruginosa* deletion mutant Δ*pslBCD*, Δ*pelA*, Δ*pilA* or Δ*fliM* were used to cultivate mixed-biofilms with the other seventeen bacterial species respectively. The Psl polysaccharide and T4P were found to be required for *P. aeruginosa* to predominate in the complex microbial biofilm community (Figure 5). Psl was particularly essential for bacterial colonization in the mixed-species biofilms (46), so Δ*pslBCD* was completely outcompeted by other species in biofilm community (Figure 5), even though its planktonic growth was similar to that of the wild-type PAO1 strain (Supplementary Figure 5).

### Quorum sensing is not required for the fitness of *P. aeruginosa* in the mixed-species biofilm community

Quorum sensing systems are well known to play important roles in biofilm formation both *in vitro* and *in vivo* (47–52). Previous studies showed that *P. aeruginosa* is able to outcompete other microbial species by releasing quorum sensing-regulated virulence factors, such as rhamnolipid, elastase, exotoxins, pyocyanin and pyoverdine (53–57). However, the expression of the *P. aeruginosa* quorum sensing-related genes, such as *lasA*, *lasB*, *rhlA*, *rhlB* and *pqsH*, were found to be downregulated in this biofilm community compared to its mono-species biofilm (Table 2), suggesting that quorum sensing is not important in contributing to the fitness of *P. aeruginosa* under this condition. Intriguingly, deleting the *lasR* or *mvfR* quorum sensing genes in *P. aeruginosa* even increased this species’ fitness over that of other species in the mixed-species biofilms compared to the PAO1 wild type (Figure 5), which might be because *P. aeruginosa* H1-T6SS for competition was repressed by LasR and MvfR (58).

## Discussion

Our study reported on the establishment of physiologically reproducible laboratory planktonic and biofilm communities and their use in investigating how *P. aeruginosa* competes with multiple other bacterial species. We tracked the population dynamics and profiled the transcriptomes of the established communities and showed that *P. aeruginosa* maintains distinct transcriptomes in complex microbial communities compared to its mono-species cultures (Figure 4). The H1-T6SS, Psl and T4P were found to be the key components for *P. aeruginosa* to gain fitness over other species in the mixed-species biofilm communities (Figure 5 and Table 2). These pathway-involved genes, such as *clpV1* (6.8-fold), *pilA* (17.1-fold) and the *pslBCD* operon, were all significantly upregulated in the microbial biofilm community (*P. aeruginosa* is the dominant species) as compared to the planktonic cultures (*P. aeruginosa* is not the dominant species). Our results highlight the biological significance of biofilm formation for *P. aeruginosa* to gain fitness in complex microbial communities (59), and we confirmed that H1-T6SS is a powerful weapon for *P. aeruginosa* interspecies competition within the ecological niche in bacterial communities (39, 41). Our findings on *P. aeruginosa* in the mixed-species microbial community are in accordance with previous studies in pure culture or two species mixture (18, 19, 26, 40) and suggest that Psl, T4P and H1-T6SS are potential targets to control or eliminate *P. aeruginosa* from complex microbial biofilm communities, such as the microbial consortia from medical devices.

**Figure 4.**
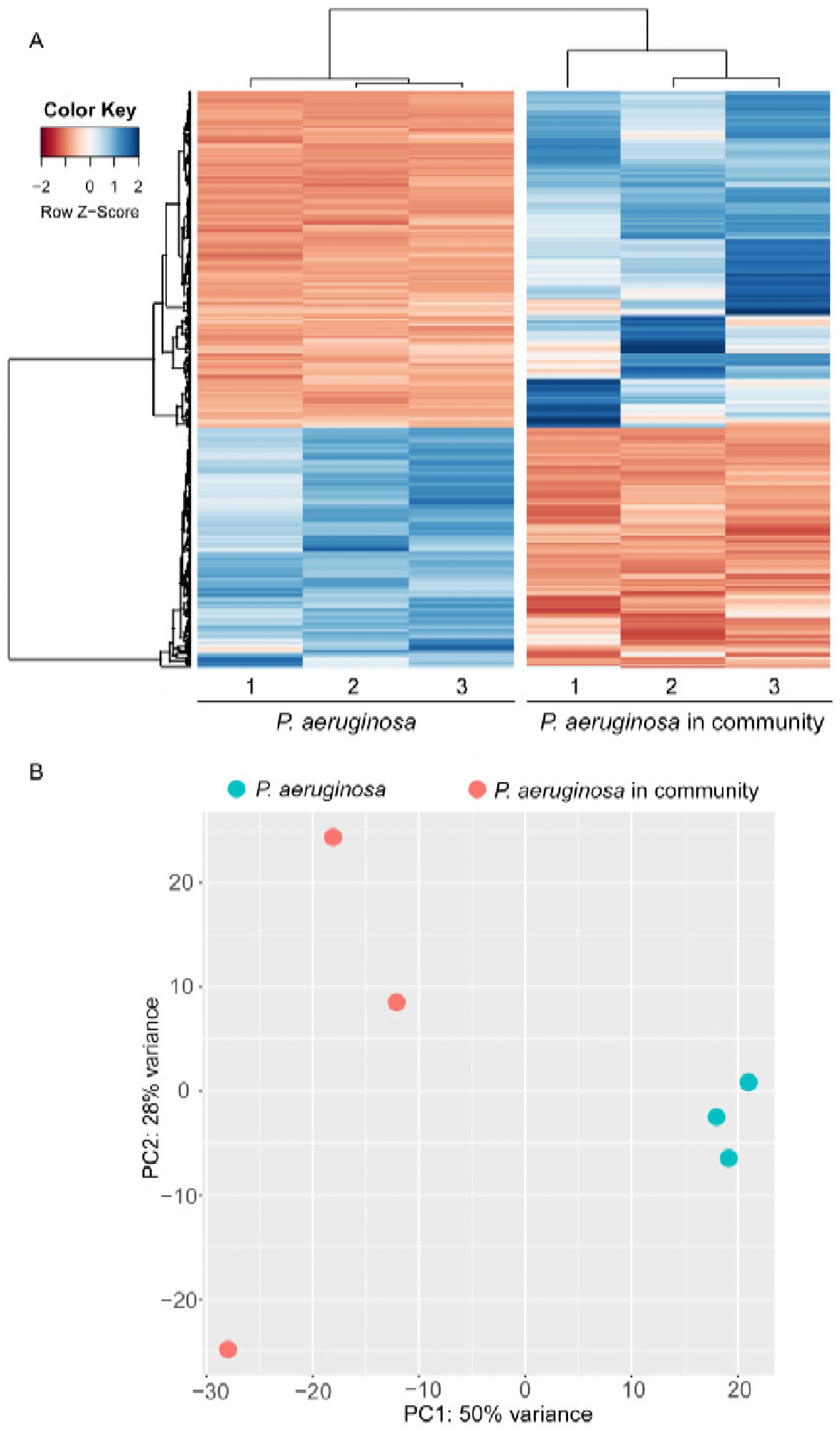
Transcriptomic comparison of *P. aeruginosa* between mono-species biofilm cells and the mixed-species biofilm community. A) A heat map representing *P. aeruginosa* genes expression level up-or downregulated more than 2-fold in the mixed-species biofilm compared to the mono-species biofilm. The Z-Score shows the standardization of each gene expression level between the two different groups. Detailed information on this regulation is listed in Supplementary Table 3. B) A principal component analysis (PCA) shows that the expression profiles of *P. aeruginosa* between the mono-species biofilm and it in the mixed-species biofilm are different. Each dot indicates one biological replicate with different colours representing the sample source.

**Figure 5.**
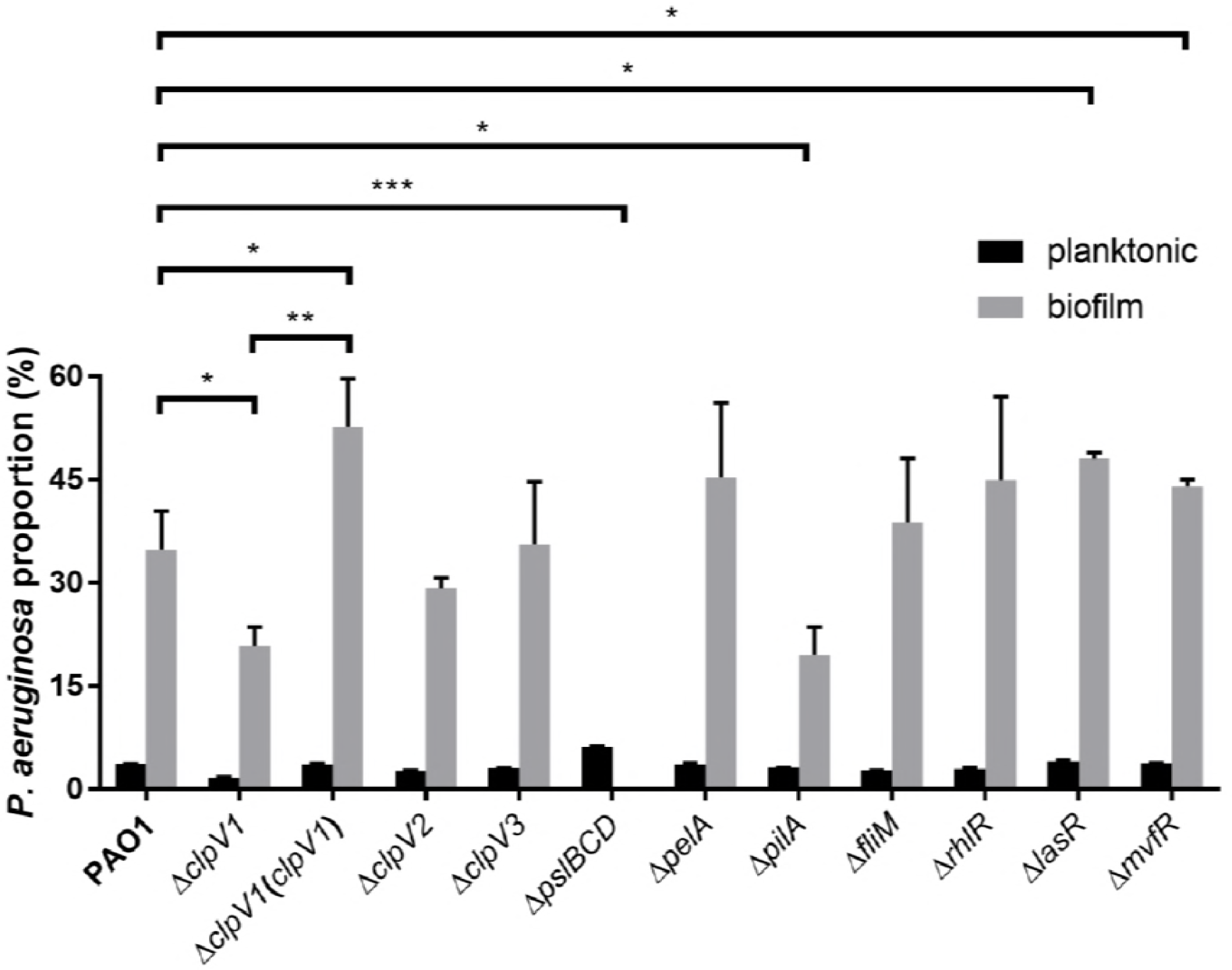
*P. aeruginosa* population dynamics in the mixed-species biofilm community affected by H1-T6SS and biofilm formation determinants. Each column shows the population dynamics of GFP-tagged *P. aeruginosa* parental strain PAO1 and the knockout mutants in the mixed-species microbial planktonic (black) and biofilm (grey) communities at different heights with the standard deviation of three biological replicates. The asterisk indicates statistical significance by t-test with p<0.05.

In addition, we demonstrated the sensitivity of the single molecule detection technique, NanoString nCounter® 16S rRNA array, in monitoring the population dynamics in complex bacterial communities. Our results showed that the accuracy of this technique depends on the specificity and affinity of the probes, which need to be normalized during the calculation of population dynamics.

*P. aeruginosa* is able to dominate airway infection microbiota (60) and wound communities after the administration of antibiotics (61), suggesting that *P. aeruginosa* has a competitive physiology in mixed-species microbial communities. Our transcriptomic analysis showed that two of the primary mechanisms for *P. aeruginosa* to maintain this competitive advantage are H1-T6SS and biofilm formation, which are different from most previous reports in which many quorum sensing-regulated virulence factors, such as elastase, PQS, pyoverdine and pyocyanin, were used by *P. aeruginosa* to kill other bacteria (53–57). The reasons for this difference may be as follows: 1) *P. aeruginosa las* system and *pqs* system repress H1-T6SS to compete with other bacteria (58); 2) quorum sensing signalling molecules and virulence factors might function as signals to induce aggressive phenotypes from bacterial competitors as well as hosts via cross-talk, which in turn arrests the growth of *P. aeruginosa;* 3) the high production level of quorum sensing-regulated virulence factors might consume a great deal of energy to reduce the replication of *P. aeruginosa* cells or even trigger autolysis phenotypes (62); or 4) the flow biofilm cultivation system with continued fresh medium flow through the biofilms might wash away the secreted quorum sensing signals and virulence factors and thus reduce the efficacy of quorum sensing. Thus, the cell-contact based T6SS might be the most efficient mechanism for *P. aeruginosa* to outcompete other species. This is supported by the data that loss of ClpV1 in H1-T6SS will decrease the competitive capacity of *P. aeruginosa* in our laboratory model (Figure 5). Recently, Allsopp *et al* has proved that all three T6SS in *P. aeruginosa* express more at 25°C than them at 37°C (63), however, expressed H2-T6SS and H3-T6SS did not affect *P. aeruginosa* competition capacity as H1-T6SS at room temperature in our experiment (Figure 5).

H1-T6SS is well known for its roles in competitiveness and pathogenicity in *P. aeruginosa* (26, 39). In addition to H1-T6SS, Psl and T4P are critical components of *P. aeruginosa* fitness in the mixed-species biofilm community (Figure 5). Our result is in accordance with previous research showing that secreted protein A from *S. aureus* can inhibit *P. aeruginosa* biofilm formation by binding to Psl and T4P (64). One limitation of our current study is that we have not elucidated whether *P. aeruginosa* employs T6SS, Psl and T4P sequentially or simultaneously when outcompeting the other species in the biofilm community. Additionally, further study is required to validate whether the T6SS, Psl and T4P are required by *P. aeruginosa* to outcompete other species *in vivo.* Because the amounts of bacteria from *in vivo* samples are usually low and insufficient for transcriptomic analysis, we can explore suitable probes for the NanoString nCounter® array to examine the population dynamics and gene expression of *P. aeruginosa* from the *in vivo* communities. Moreover, *K. pneumonia* is the secondary dominant species in the biofilm co-cultures but not in planktonic co-cultures like *P. aeruginosa* (Figure 2), and it constantly co-exists with *P. aeruginosa in vitro* and *in vivo* (65). Thus, it would be interesting to study its survival mechanism in the future.

## Methods

### Bacterial strains, media and growth conditions

The bacterial species used in this study are listed in Supplementary Table 1. All these strains were revived in tryptic soy broth (TSB) agar plates and subsequently in TSB liquid medium at 30°C. Mixed-species communities were cultivated in 10% TSB at room temperature. To construct *P. aeruginosa* gene knockout mutants and green fluorescent protein (GFP)-tagged strains, AB minimal medium (66) supplemented with 10 mM citric acid supplemented with appropriate antibiotics was used to select *P. aeruginosa. E. coli* was grown in LB medium at 37°C. 60 μg/ml of gentamicin was used to select *P. aeruginosa,* while 15 μg/ml of gentamicin, 10 μg/ml of chloramphenicol and 100 μg/ml of ampicillin were used to select *E. coli,* when appropriate.

### Growth assay and static biofilm cultivation

Overnight cultures of each bacterial species were diluted to OD_600_ of 0.01 in 10% TSB, 150 μl diluted culture were loaded into 96-well microtitre plates. The plates were incubated in an infinite M200PRO multimode microplate reader (Tecan Schweiz AG, Männedorf, Switzerland) at room temperature, and the OD_600_ of each well was measured and recorded hourly. A 10% TSB blank medium was used as a negative control, and its reading has been subtracted from all the culture readings to plot the growth curves for each species.

To examine the static biofilm-forming capacities of each species, bacterial cultures were prepared using the same procedures as mentioned in growth assay. After overnight cultivation, the planktonic cells were discarded from the microtitre plate, and the wells were washed three times using tap water to remove any residual planktonic cells. A 180 μl volume of 0.1% crystal violet (CV) solution was loaded into each well to stain the biofilm for 15 min. The excess stain in the wells was washed out with tap water three times. Finally, 180 μl of 30% acetate acid was loaded into each well to dissolve CV from the stained biofilm. The optical intensity at OD_550_ was measured by an Infinite M200PRO multimode microplate reader to quantify the biofilms formed by each bacterial species.

### Cultivation of the mixed-species planktonic and biofilm communities

Overnight cultures of the eighteen individual bacterial species were diluted in 10% TSB medium and normalized to the same ratio into the primary mixture where OD_600_ = 1. The primary mixture was then diluted to OD_600_=0.01 in 10% TSB, and this diluent was used as the initial inoculum for both planktonic cultures in shake flask and biofilm cultures in flow tube reactors. The 24-hour-old planktonic cultures and 5-day-old biofilm cultures were harvested for population dynamic or transcriptomic analysis.

### RNA preparation for sequencing

The bacterial cells were first treated with RNA Protect Reagent (Qiagen®, Germany) to maintain the integrity of RNA. The total RNA was extracted from these bacterial cells using a miRNeasy Mini Kit (Qiagen®, Germany) with modifications. A Turbo DNA-free Kit (Thermo Fisher Scientific®, Lithuania) was used to remove genomic DNA contaminant from total RNA. DNA contamination was assessed with a Qubit®dsDNA High Sensitivity (HS) assay (PicoGreen dye) and a Qubit® 2.0 Fluorometer (Invitrogen®, Austria) according to manufacturer’s instructions. Ribosomal RNA was depleted with a Ribo-Zero rRNA removal Kit (Illumina, USA). The integrity of the total RNA was assessed with an Agilent TapeStation System (Agilent Technologies, UK).

The Double-stranded cDNAs were reverse-transcribed using a NEBNext RNA first and second strand synthesis module (NEB®, USA). cDNAs were subjected to Illumina’s TruSeq Stranded mRNA protocol. The quantitated libraries were then pooled at equimolar concentrations and sequenced on an Illumina HiSeq2500 sequencer in rapid mode at a read-length of 100 bp paired-ends.

### NanoString nCounter® population analysis

One ng of purified total RNA from the different bacterial cultures was analysed using a NanoString nCounter® Gene Expression CodeSets platform (NanoString Technologies, Inc., United States) with the customized CodeSets for our selected bacterial species (Supplementary Table 1) according to the manufacturer’s protocol to obtain raw counts. The quality control assessment and raw counts were normalized and analysed using nSolver™ analysis software version 2.6 (NanoString Technologies, Inc., United States). The samples were subjected to manufacturer-recommended default parameters. The geometric mean was selected for negative control subtraction, while the geometric mean was used to compute the normalized factor for positive control normalization and the flag lane if the normalization factors were outside the 0.3 – 3 range. The experiments were performed in triplicates, and the results are presented as the means ± S.D.

### Transcriptomic analysis

The accession number for RNA-seq is SRP128411. For *P. aeruginosa* transcriptomics analysis, PAO1 genome (NC_002516) was used as the reference with annotation from the Pseudomonas Genome Database (www.pseudomonas.com). The RNA-Seq raw data were analysed using the “RNA-Seq and expression analysis” application in CLC genomics Workbench 10.0 (QIAGEN). The total gene reads from CLC genomics Workbench 10.0 were subjected to the DESeq2 package for statistical analysis (67) by using R/Bioconductor (68). A hierarchical clustering analysis was performed with a negative binomial test using the DESeq2 package. The heatmap.2 package was used to draw a heat map for the differentially expressed genes of *P. aeruginosa* cells with fold change larger than 2 and an adjusted p-value smaller than 0.05. And the principal component analysis (PCA) plot was generated in R/Bioconductor.

The RNA sequences adapter-trimmed and assembled RNA sequences of the mixed-species microbial community were used as metatranscriptomics data for analysis. The sequences of the mixed-species communities grown as biofilm and planktonic cells were aligned against the NCBI non-redundant (NR) protein database using DIAMOND with default settings. The output aligned gene reads in DAA format were uploaded and meganized in MEGAN6.11.1 with a minimum bit-score of 50 and a top percentage of 25 (69). Functional analyses of these aligned gene reads were performed using SEED classifications. A Principle Coordinates Analysis (PCoA) was plotted to cluster the samples based on a functional analysis. The functional analysis results are illustrated in stacked bar charts.

### Fluorescence-based *P. aeruginosa* population assay

To develop a rapid *P. aeruginosa* population dynamics assay in the mixed-species cultures, a single copy of a green fluorescent protein (GFP) tag was inserted into the *P. aeruginosa* chromosome by using a mini-Tn7-Gm-gfp transposon (70). The other seventeen bacterial species without *P. aeruginosa* were used as a background control. The GFP fluorescence readings of both planktonic and biofilm cultures, with gradient proportion of GFP-tagged *P. aeruginosa* derived strains, were recorded using a Tecan infinite M200PRO microplate reader. Then the fluorescence-proportion standard curves of each tested strain were generated from these GFP readings (Supplementary Figure 4). The proportions of *P. aeruginosa* derived strains in communities can be calculated using these fluorescence-proportion standard curves.

### Constructions of *P. aeruginosa* gene deletion and complementary mutants

The upstream fragment of target gene was amplified by primer-1 and primer-2, the downstream fragment of the target gene was amplified by primer-3 and primer-4 (Supplementary Table 4), and then both fragments were fused into a pK18-Gm-mobsacB plasmid (71) and transformed into the *P. aeruginosa* PAO1 wild-type strain to delete the target gene as previously described (71). PCR and sequencing were used to confirm the selected mutant strains (Supplementary Table 5).

To complement the *P. aeruginosa clpV1* mutant, a DNA fragment from the *clpV1* upstream 22 bps to the downstream 31 bps was amplified by primers *clpV1-up* and clpV1-down (Supplementary Table 4) and double-digested by *BamHI* and *HindIII.* The purified fragment was ligated with the pUCP22 vector (72), which was digested by the same restriction enzymes to construct the pUCP22-clpV1 complementary plasmid. After verification by sequencing, the pUCP22-clpV1 plasmid was transformed into the *P. aeruginosa* Δ*clpV1* mutant by electroporation to construct a *clpV1* complementary strain.

### Confocal microscopy imaging

*P. aeruginosa* PAO1-GFP-containing planktonic cultures and biofilm cultures (the same samples as those used for the fluorescence based population assay) were stained with 5 uM SYTO^TM^ 62 Red fluorescent nucleic acid stain (Molecular probe) to stain all cells for 15 min before 5 ul of each sample was aliquot onto the glass slide and covered with cover slip. The samples were visualized and confocal images were acquired using confocal laser scanning microscopy (CLSM) Zeiss LSM 780 (Carl Zeiss, Germany) with a 63x 1.40 DICII objective. Green and red fluorescence were observed upon 488 nm and 633 nm excitation wavelengths, respectively.

## Acknowledgements

This research was supported by the National Research Foundation and Ministry of Education
Singapore under its Research Centre of Excellence Program and AcRF Tier 2 (MOE2016-T2-1-010) from Ministry of Education, Singapore. We appreciate Professor Zhang Lian-Hui to provide us the pK18-Gm-mobsacB plasmid. We thank Dr. Daniela Moses, Mr. Yap Zhei Hwee, Rikky Wenang Purbojati and Mr. Muhammad Hafiz Bin Ismail for help with the RNA-seq experiments.

## Author contributions

Y. C. designed and carried out experiments, interpreted results and wrote the manuscript. L. Y. designed the project and revised the manuscript. J. K. H. Y. helped with the confocal microscopy imaging and biofilm bioreactors. Z. C. performed metatranscriptomic analysis. Y. D., L. Z. and Y. D. helped to revise the manuscript.

## Electronic supplementary material

Supplementary information is available at Applied and Environmental Microbiology’s website.

